# Collective forces of tumor spheroids in three-dimensional biopolymer networks

**DOI:** 10.1101/654079

**Authors:** Christoph Mark, Thomas J. Grundy, David Böhringer, Julian Steinwachs, Geraldine M. O’Neill, Ben Fabry

## Abstract

We describe a method for quantifying the contractile forces that tumor spheroids collectively exert on highly nonlinear three-dimensional collagen networks. While three-dimensional traction force microscopy for single cells in a nonlinear matrix is computationally complex due to the variable cell shape, here we exploit the spherical symmetry of tumor spheroids to derive a scale-invariant relationship between spheroid contractility and the surrounding matrix deformations. This relationship allows us to directly translate the magnitude of matrix deformations to the total contractility of arbitrarily sized spheroids. We show that collective forces of tumor spheroids reflect the contractility of individual cells for up to 1h after seeding, while collective forces on longer time-scales are guided by mechanical feedback from the extracellular matrix.

## Main

In the pathological process of tumor invasion, cancer cells leave a primary tumor, either individually or collectively^1^. This process requires that cells exert physical forces onto the surrounding extracellular matrix^2, 3^. Numerous biophysical assays have been developed to quantify the traction forces of individual cancer cells by measuring the deformations that a cell induces in linear elastic substrates (2D and 3D) with known stiffness^4–6^. To mimic the physiological condition of cells invading connective tissue in vitro, however, cells are typically seeded into non-linear biopolymer networks such as reconstituted collagen, which stiffens significantly when extended^7, 8^ but softens when compressed^8, 9^. Considering these nonlinear material properties in a finite element approach allows for the reconstruction of the three-dimensional traction force field around individual cells in a biopolymer network^9^.

Here, we extend this method to make it applicable to multicellular aggregates (so-called spheroids) consisting of several hundreds or thousands of individual cancer cells that are embedded in a collagen gel (Fig. 1a-c). The motivation for this extension is that tumor spheroids are able to replicate the main structural and functional properties of solid tumors^10^, including the individual and collective invasion into the surrounding matrix^11^. In particular, it has been hypothesized that the collective traction force of a whole tumor may alter the mechanics of the extracellular matrix and facilitate invasion by aligning collagen fibers radially away from the tumor^12–14^. However, current 3D finite element force reconstruction methods for single cells, especially in a non-linear material such as collagen, are computationally too slow for analyzing large (~0.5 mm) tumor spheroids. Moreover, measurements typically require a confocal microscope equipped with a high-resolution (NA 1.0 or higher) water dip-in long working distance objective to image the three-dimensional structure of the collagen fiber network using reflection microscopy.

**Figure 1.**
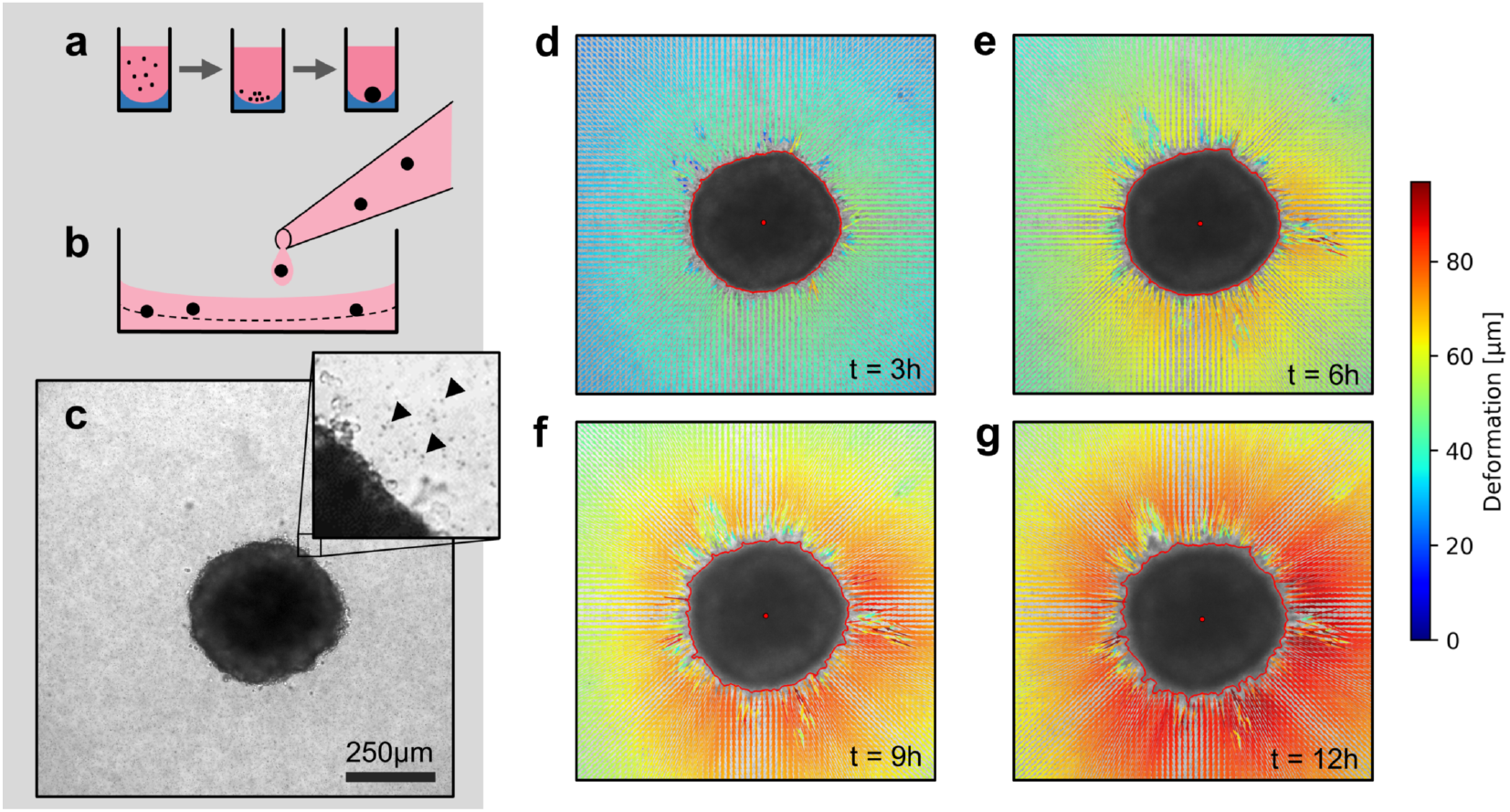
Spheroid formation and collagen contractility assay. **a:** Spheroid generation process within agarose-coated 96-well plates. **b:** Spheroid embedding process in collagen gels. The spheroids are suspended in a collagen solution and subsequently pipetted onto a pre-poured layer of collagen (indicated by the dashed line). **c:** Exemplary brightfield image of the equatorial plane of a U87 spheroid containing 7,500 cells. The inset shows the edge of the tumor spheroid and the micron-sized fiducial markers (arrows) that are added to the collagen solution. **d-g:** Deformation field obtained by Particle Image Velocimetry, 3 h, 6 h, 9 h and 12 h after the collagen gel has polymerized. The spheroid outline is determined by image segmentation and indicated by the red line.

To overcome these technical challenges, we forgo subcellular force resolution and exploit the approximately spherical symmetry of tumor spheroids (Supplementary Fig. 1). Accordingly, it is sufficient to measure the far-field deformations of the surrounding collagen matrix from a single slice through the equatorial plane of the spheroid, eliminating the need for high-resolution 3D imaging. Image acquisition can be done with low resolution (4x-10x objective, NA 0.1) brightfield microscopy in combination with micron-sized fiducial markers embedded in the collagen gel to quantify matrix deformations over time (Fig. 1c).

In this report, we track the deformations of the collagen matrix induced by the contractile forces of A172 and U-87 MG (referred to as U87 hereafter) glioblastoma spheroids from brightfield time-lapse images (every 5 min) using Particle Image Velocimetry^15^. In general, we find that the spheroids induce an approximately radially symmetric, inward-directed deformation field with monotonically increasing absolute deformations over time (Fig. 1d-g; Supplementary Videos 1,2), in line with a previous report on CT26 colon carcinoma cells^16^.

To relate the measured deformation field surrounding a spheroid to physical forces generated by the cells, we use the finite element approach described in Ref. 9. Specifically, we simulate a small spherical inclusion with a negative hydrostatic pressure (that emulates contracting cells within the inclusion) within a large surrounding volume of collagen (Fig. 2a,b). This computational analysis predicts that the absolute deformations of the collagen *u*(*r*) are largest directly at the boundary of the inclusion and fall off with increasing distance *r* from the center, depending on the pressure (Fig. 2b). For a given pressure, the absolute deformations increase with the radius *r*_0_ of the inclusion. Importantly, when normalized by the radius of the inclusion *r*_0_, the deformations *u*/*r*_0_ collapse onto a single curve when plotted against the normalized distance *r*/*r*_0_ (Fig. 2c, Supplementary Note 1). This implies that the shape of the simulated deformation field only depends on the pressure but not on the size of the inclusion (i.e. on the spheroid radius *r*_0_ at the time of seeding).

**Figure 2.**
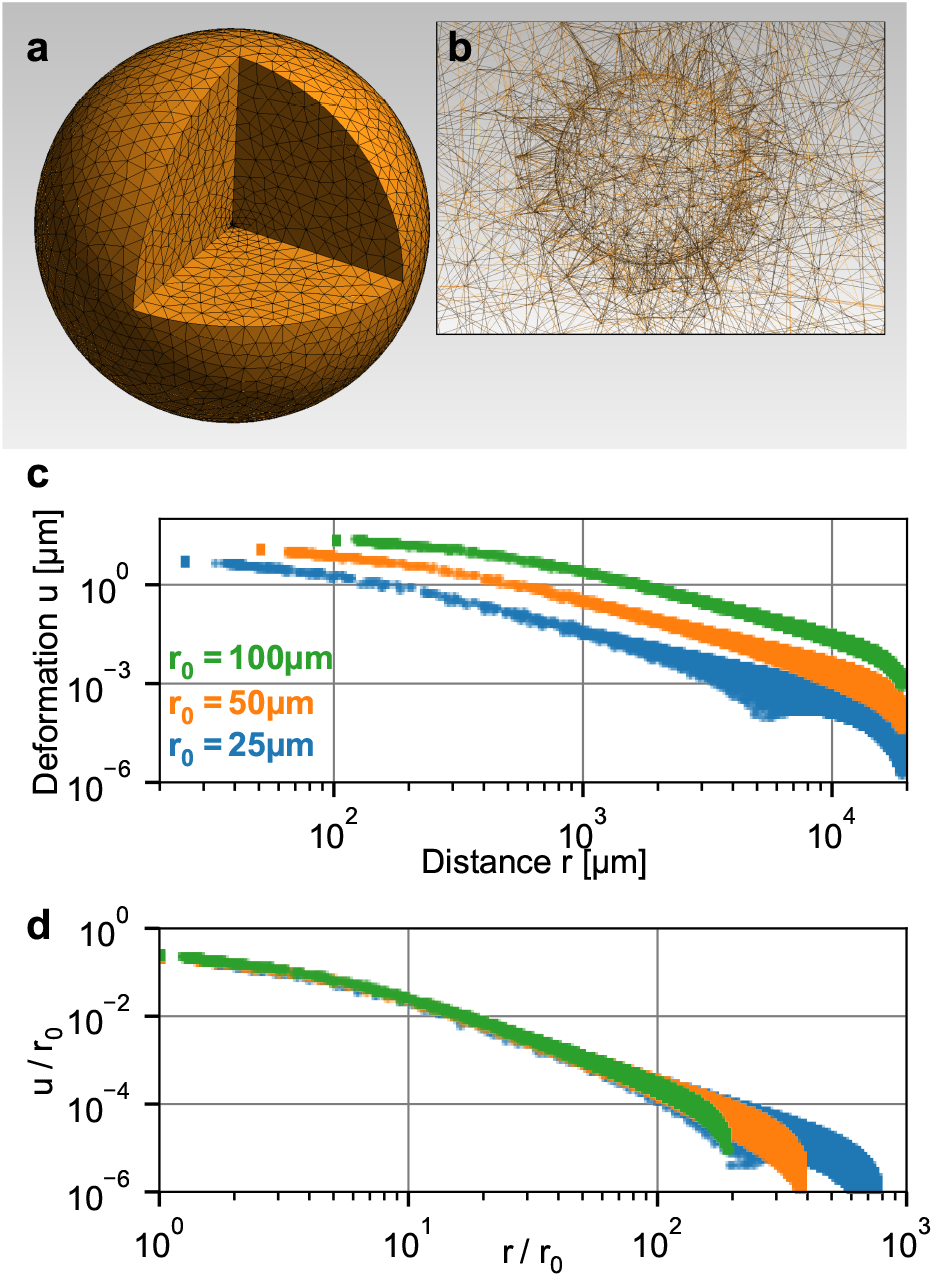
Simulation of a spherical inclusion in collagen. **a:** Illustration of the tetrahedral mesh used for the material simulation. The spherical volume has a radius of 2 cm, with a spherical inclusion in the center. **b:** Enlarged section of the tetrahedral mesh around the spherical inclusion with a radius of *r*_0_ = 100 μm. **c:** Simulated absolute deformations 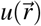 as a function of the distance 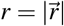 from the center of the volume, for an inward-directed pressure of 100 Pa acting on the surface of the inclusion. Different colors indicate different radii *r*_0_ of the spherical inclusion. **d:** Same as in (c), but with deformations and distances normalized by *r*_0_. For a given inbound pressure, all curves collapse onto a single relationship.

The collapse of the normalized deformation versus distance relationship furthermore implies that we can estimate the contractile pressure of a tumor spheroid of arbitrary size from a look-up table. To create this look-up table, we perform 120 simulations with pressures ranging from 0.1 Pa to 1000 Pa. The simulated deformation fields are normalized by *r*_0_, binned and interpolated to obtain smooth deformation curves (Fig. 3a). For a low pressure of ~1 Pa, the deformation field as a function of radial distance from the spheroid center falls off with increasing distance according to a power law with an exponent (= slope in a double logarithmic plot) of *α* = −2, as expected for a linear elastic material. With increasing pressure, however, the deformations near the spheroid surface fall off more slowly, with a slope approaching values around *α* = −0.2 for high pressure values >1000 Pa (Supplementary Fig. 2), indicating long-range force transmission due to a stiffening of the collagen fibers. This is in line with reported theoretical models^17, 18^ and experimental findings^19^. When we overlay predicted and measured deformation curves around tumor spheroids, we find good agreement in particular for high forces (Fig. 3a), confirming the model assumptions and the computational approach (Supplementary Fig. 3).

**Figure 3.**
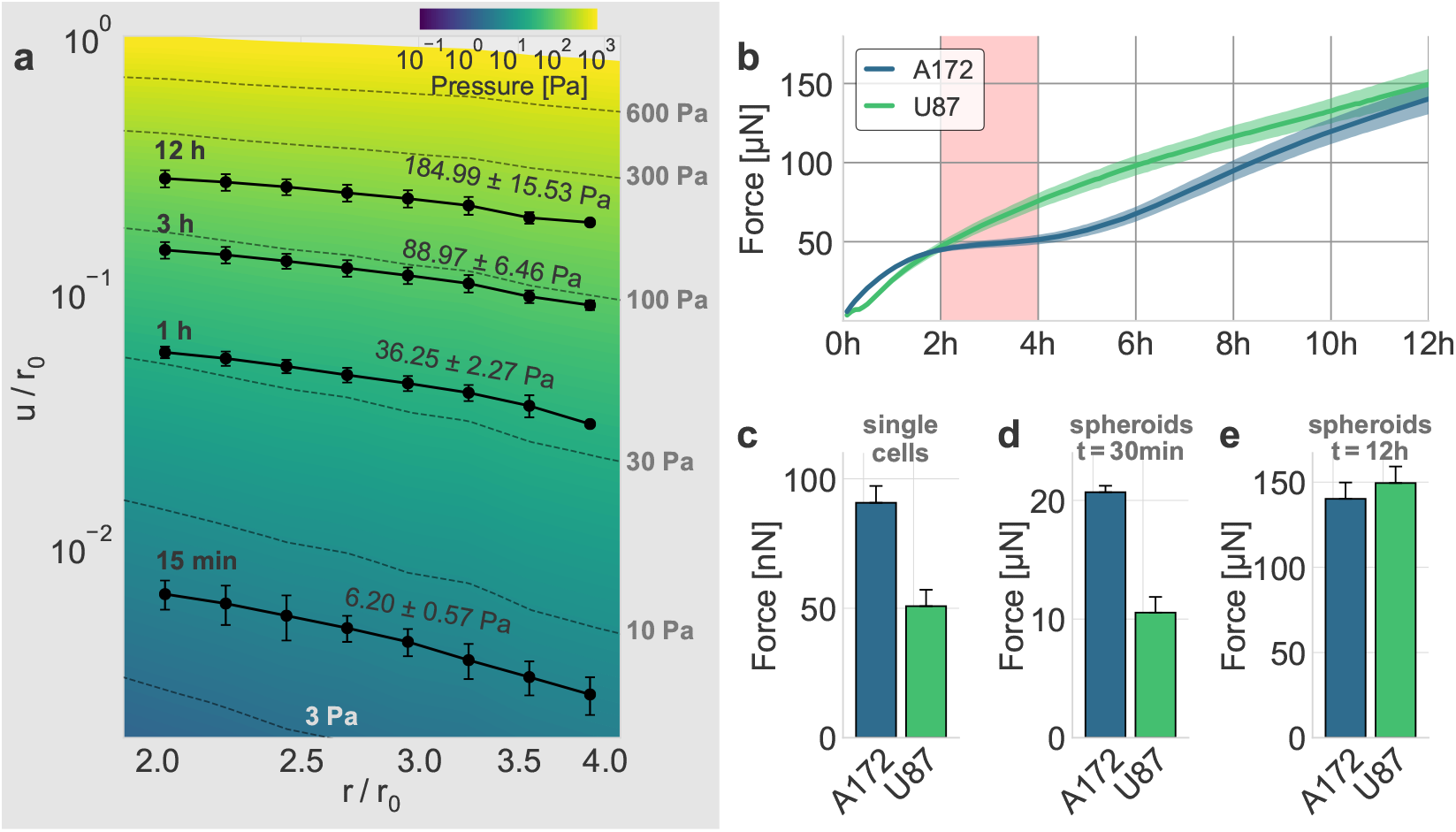
Collective contractility of glioblastoma spheroids. **a:** Normalized deformations as a function of the normalized distance for material simulations of varying pressure (color coding & dashed lines). Solid black symbols show the measured strain in the collagen matrix at different time steps for an exemplary U87 spheroid. The best-fit pressure values are noted next to the strain curves. **b:** Time course of the mean contractility and corresponding standard error (shaded) for A172 (blue) and U87 (green) spheroids. The 2 h-resting period of the A172 spheroids is marked in red. **c:** Median cell contractility as measured by single-cell 3D traction force microscopy (A172: n=90; U87: n=86). **d:** Mean collective cell contractility of tumor spheroids after 30 min measurement time. **e:** Mean collective cell contractility of tumor spheroids after 12 h measurement time. Error bars denote 1 standard error.

We apply this assay to two glioblastoma cell lines, A172 (15,000 cells per spheroid) and U87 (7,500 cells per spheroid, to match the size of A172 spheroids; Supplementary Fig. 1). Although collagen is present in only small amounts in the normal human brain, glioblastoma cells readily bind to collagen^20^. Moreover, recent studies have shown that fibrillar collagens are an integral part of the locally produced extracellular matrix in glioblastomas^21, 22^, yet little is known about the traction forces exerted by GBM in order to facilitate movement through brain tissue. Furthermore, collagen is found in the basement membrane surrounding blood vessels, which are a major route of GBM invasion^20, 23^. Reconstituted collagen gels display a Young’s modulus of 162 ± 25 Pa^9^ in the linear regime, closely emulating the soft environment of the brain (100-1000 Pa^24^).

We find that with increasing time after seeding, U87 spheroids monotonously increase their contractility (pressure × surface area; Fig. 3b, Supplementary Fig. 4) over the 12 h observation period. By contrast, A172 spheroids consistently show a 2 h-resting period after a fast initial increase in contractility. While we have observed temporal variations in the contractility of individual breast carcinoma cells in a previous study^9^, such a collective resting phase requires a synchronized change in cellular force generation across the whole cell population. A likely mediator for this cell-cell coupling is the collagen matrix: as the cells pull on the matrix, collagen exhibits strain stiffening. This change in material stiffness then provides a mechanical feedback to the cells and may thus alter cell behavior at the population level.

When we compare the collective contractility of tumor spheroids to the contractility of individual cells of the respective cell lines (Supplementary Fig. 5), we find that after 12 h, A172 spheroids and U87 spheroids of equal size attain comparable collective contractility of 140 ± 10 μN and 149 ± 10 μN (Fig. 3e). By contrast, single-cell 3D traction force microscopy indicates that individual A172 cells are nearly 2-fold stronger compared to U87 cells (91 nN vs. 51 nN) (Fig. 3c). Thus, collective contractility is not necessarily related to the respective traction forces of individual cells. However, collective contractility of A172 spheroids and U87 spheroids observed at an early time point (30 min after seeding) exactly reflect the differences seen at an individual cell level: A172 spheroid contractility is increased by 96% compared to equally sized U87 spheroids (Fig. 3d). During these initial time steps, the induced strains on the collagen matrix are still small, and hence there is no global stiffening of the material which feeds back to cell behavior.

We conclude that our contractility assay is able to quantify the collective force generation process in tumor spheroids containing hundreds or thousands of cells. For A172 and U87 glioblastoma cells, we find that the collective forces reflect the contractility of individual cells during the initial contraction phase (≲1 h), but not on longer time scales. In particular, the large strains induced by the spheroids may significantly alter the mechanical environment of the invading cells due to strain stiffening and fiber alignment, and thus alter cellular force generation at a collective level and potentially induce enhanced invasion into the surrounding tissue. As cell-matrix interactions and the process of tissue remodelling are increasingly recognized as therapeutic targets^25^, our method provides a reliable and simple in-vitro assay to quantify the mechanics behind collective effects in cancer invasion that cannot be measured on a single-cell level.

## Acknowledgements

This work was supported by Deutsche Forschungsgemeinschaft grants FA-336/11-1, STR 923/6-1, the Research Training Group 1962 “Dynamic Interactions at Biological Membranes: From Single Molecules to Tissue”, the German Academic Exchange Service (DAAD) project “Physical mechanisms in glioblastoma cell invasion”, and National Institutes of Health grant HL120839. T.G. was supported by an Australian Government Research Training Program Scholarship and generous funding from the Petersen Foundation.

## Author contributions

B.F. and G.ON. designed the study. C.M., D.B., and J.S. developed the force reconstruction method. T.G. performed the experiments. C.M. and T.G. performed the analyses. C.M., T.G., G.ON., and B.F. wrote the paper. All authors read and approved the final manuscript.

## Competing interests

The authors declare no competing interests.

## Methods

### Cell culture

A172 and U-87 MG (referred to as U87 in the main text) glioblastoma cell lines are cultured at 37°C, 95% humidity and 5% CO_2_ in DMEM High Glucose Pyruvate with 10% (volume/volume) fetal bovine serum, and 100 Units/ml penicillin/streptomycin (all Thermo Fisher Scientific). Cell lines are short tandem repeat (STR) profiled to confirm identity (CellBank Australia) and are confirmed negative for mycoplasma contamination with Venor GeM Classic detection kit (Minerva biolabs).

### Spheroid culture

Spheroids are created from low-adherent, concave-bottomed surfaces in 96-well dishes^26^. 50 μl of a heated 1.5% (weight/volume) agarose (Thermo Fisher Scientific)/DMEM gel solution is pipetted into the wells of a 96-well dish. Following a 10-15 min interval, the solution cools and forms a non-adherent, concave surface.

Subsequently, cells are detached from their tissue culture flasks with 0.05% trypsin solution, counted (15,000 cells per dish for A172 cells and 7,500 cells per dish for U87 cells) and pipetted into wells containing 100 μl cell culture medium. The agarose surface promotes formation of a single spheroid per well. Spheroids take 3 days to fully form while being incubated at standard TC conditions.

### Collagen synthesis

Collagen gels are synthesized as described in Ref. 9 and consist of a 1:1 mixture of rat tail collagen (Collagen R, 2 mg/ml, Matrix Bioscience, Berlin, Germany) and bovine skin collagen (Collagen G, 4 mg/ml, Matrix Bioscience), plus 10% (vol/vol) NaHCO_3_ (23 mg/ml) and 10% (vol/vol) 10 × DMEM (Gibco). The pH of the solution is adjusted to 10 with 1 M NaOH. For a collagen concentration of 1.2 mg/ml, the solution is diluted with a mixture of 1 volume part NaHCO_3_, 1 part 10 × DMEM and 8 parts H_2_O, at a ratio of 1:1.

### Spheroid embedding

FluoSphere polystyrene beads (1 μm diameter, Thermo Fisher Scientific) are carefully suspended, without forming bubbles, in 1.2 mg/ml collagen solution at a concentration of 2·10^8^ beads/ml. 1.5 ml of this mixture is poured into a 35 mm plastic culture dish and is allowed to settle for 2.5 min at 23 °C, during which time the spheroids are prepared for embedding. The 2.5 min waiting time is too short for a full polymerization of the collagen solution but is sufficient to ensure that spheroids do not sink to the base of the dish.

After the preparation of the bottom collagen layer, 4 to 5 individual spheroids are removed from their culture plate wells and carefully transferred into a 15 ml centrifuge tube using a P1000 pipette. Once the spheroids have settled to the base of the tube, excess media is aspirated away and spheroids are gently resuspended in 500 μl of the 1.2 mg/ml collagen/bead mixture. The mixture, complete with suspended spheroids, is then transferred from the tube into the 35 mm dish using a P1000 pipette. By introducing the collagen into the dish dropwise, the positioning of the spheroids within the gel can be controlled. Spheroids are kept separate from each other and away from culture dish margins or air bubbles. After spheroid seeding, the gel is incubated at 37 °C and 5% CO_2_ for 1 h to fully polymerize. 1.5 ml of prewarmed cell media is added to the dish, and imaging is started.

### Time-lapse imaging

The equatorial plane of the embedded spheroids is imaged in brightfield mode with a 5× magnification 0.1 NA objective and a CCD camera (corresponding to a pixel size of 1.29 μm) for at least 12 h, with a time interval of 5 min between consecutive images. Samples are kept in a stage-mounted incubation chamber (37 °C, 5% CO_2_) during time-lapse imaging. In total, we imaged 17 A172 spheroids with 15,000 cells, 14 U87 spheroids with 15,000 cells, and 13 U87 spheroids with 7,500 cells (at least three independent experiments per condition).

### Material simulations

We use the semi-affine material model described in Ref. 9 to simulate the non-linear behavior of collagen. In particular, collagen gels exhibit three different mechanical regimes, depending on the applied strain. Individual fibers buckle easily under compression (with exponentially suppressed stiffness) and only attain a constant stiffness for small strains, while they exponentially stiffen under large strains:

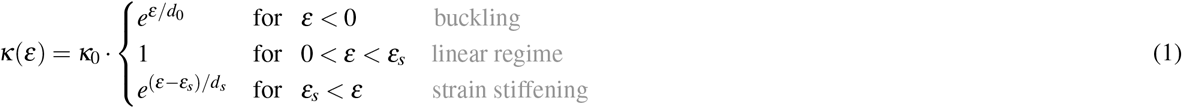

where *κ*_0_ denotes the linear stiffness, *d*_0_ and *d*_*s*_ describe the rate of stiffness variation during buckling and stiffening, respectively, and *ε*_*s*_ denotes the onset of strain stiffening.

These four parameters can be characterized by shear rheometry and by measuring the vertical contraction of a collagen gel under uniaxial stretch. In this study, we use the material parameters determined in Ref. 9, for a 1.2 mg/ml collagen solution based on a 1:1 mixture of rat tail collagen and bovine skin collagen:

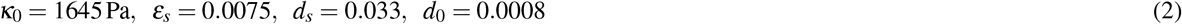

Deformations in the collagen matrix in response to inward-directed tractions at the spheroid surface are computed using a finite element approach^9^. In brief, the material volume is divided into finite tetrahedral elements, each of which is assumed to contain a number of randomly oriented fibers. When such a tetrahedron is deformed, the internal stress is first calculated by taking into account the different deformations of the contained fibers, and subsequently averaged over the faces of the tetrahedron and thus propagated to neighboring elements.

Here, we simulate a spherical bulk of material (with an outer radius of 2 cm) with a small spherical inclusion in its center (with a radius of 100 μm). The finite element mesh for this geometry is created using the open-source software Gmsh^27^. To emulate the contractile behavior of a spheroid, we assume a constant inbound pressure on the surface of the spherical inclusion and further assume zero deformations on the outer boundary of the bulk. Given these boundary conditions, we use the open-source Semi-Affine Elastic Network Optimizer (SAENO^9^) to obtain the corresponding deformation field.

### Particle image velocimetry

Given a series of images through the equatorial plane of the spheroid, we apply the open-source PIV software (OpenPIV^15^) to each pair of subsequent images. The tool then breaks up the image recorded at time *t* into *N* quadratic tiles at positions 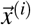 with *i* = 1, 2, …, *N* and performs a cross-correlation-based template-matching to determine the most likely offset 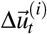 of all tiles with respect to the previous image recorded at time *t* − 5 min. These offsets represent the deformation of the material within the five minutes between two subsequent images. To account for a drift of the microscope stage between two images, we subtract the mean value

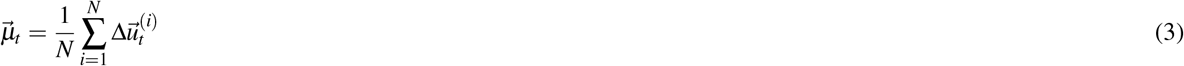

from all offsets for a given time step. To obtain the accumulated deformation 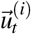 at position 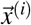 and time step *t*, we sum up the pair-wise deformation fields of all time steps *t*′ ≤ *t*:

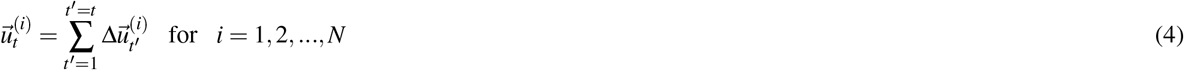

Additionally, we determine the spheroid’s centroid 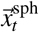 for all time steps and its initial radius *r*_0_ by image segmentation (using Otsu’s method28). As we are only interested in the radially aligned deformations towards the contracting spheroid, we compute the absolute deformations 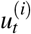 by projecting the accumulated vectorial deformations 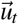 in the direction towards the spheroid center, using the relative coordinates 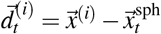:

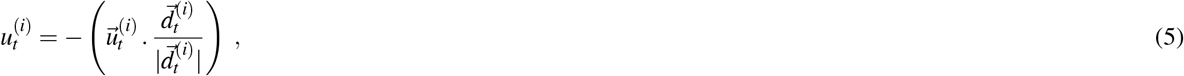

where (.) denotes the dot product. Finally, we compute the normalized deformations 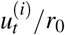 and distances 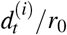 that we can directly compare to experimentally measured and normalized deformation fields.

### Force reconstruction

To assign a contractility value to a measured deformation field, we first conduct 120 material simulations assuming an inbound pressure on the surface of the spherical inclusion ranging from 0.1 Pa to 1000 Pa (logarithmically spaced) to create a look-up table. For each measured deformation vector (projected towards the center of the spheroid), we then assign the best-fit material simulation and thus a pressure value (using linear interpolation between two adjacent simulations). Finally, we take the mean of all assigned pressure values and multiply this mean value with the surface area of the spheroid (determined at the beginning of the experiment by image segmentation) to obtain the contractility.

### Single-cell 3D traction force microscopy

3D traction force microscopy is conducted as explained in Ref. 29. In brief, we pipet 1.75 ml of collagen solution into a 35 mm Petri dish and let it set for 2.5 min at room temperature. Subsequently, we suspend 15,000 cells in another 250 μl of collagen and add this solution on top to obtain a 2 mm-thick layer of collagen. This two-layer approach prevents cells from sinking to the bottom before the gel polymerizes. After waiting for one hour to ensure the complete polymerization of the gel, 2 ml of cell culture medium are added. An additional waiting time of at least two hours before imaging ensures that cells have properly spread into a polarized shape within the collagen gel. In each independent experiment, we image a cubic volume V=(370 μm)^3^ around up to 20 individual cells using confocal reflection microscopy (20× water dip-in objective with NA 1.0). We subsequently add CytochalasinD (20 μM), wait 30 min to ensure actin fiber depolymerisation and repeat the imaging. Based on the measured deformation fields, we obtain the cell contractility and force polarity of 90 individual A172 cells and 86 individual U87 cells from three independent experiments each.

### Code Availability

The traction force microscopy method introduced in this work is implemented in the Python package jointforces, which provides interfaces to both the meshing software Gmsh^27^ and the network optimizer SAENO^9^, and includes Particle Image Velocimetry functions to analyze time-lapse image data. The software is open source (under the MIT License) and is hosted on GitHub (https://github.com/christophmark/jointforces). The figures in the work have been created using the Python packages Matplotlib^30^ and Pylustrator^31^.

### Data Availability

Apart from the exemplary data that accompanies the open-source software package introduced in this work, the datasets analysed in this study are available from the corresponding author on request.

