## Supplementary Information for "Collective forces of tumor spheroids in three-dimensional biopolymer networks"

### Supplementary Note 1: Geometrical scaling in nonlinear elastic materials

The scale-invariance of the deformation field of a contracting spherical inclusion within a large body of non-linear elastic material (shown by simulation in Fig. 2 in the main text) can be derived analytically. The following derivation is discussed in more detail in Supp. Ref. 1.

Given a displacement field $\vec{U}(\vec{r})$ of an equilibrium configuration (e.g. induced by a spheroid with a radius of 100 µm and an inbound pressure of 50 Pa), we define the re-scaled displacement field (e.g. induced by a spheroid with a radius of 200 µm and the same pressure of 50 Pa) as

$$\vec{U}^*(\vec{r}) = a \cdot \vec{U}(\vec{r}/a) \tag{1}$$

with a being the scaling factor ($a = 2$ for the exemplary spheroid sizes noted above). To check whether mechanics remain unaltered by re-scaling the displacement field in this way, we need to show that the deformation gradient $\underline{F}(\vec{r})$ and thus the strain energy density $W(\vec{r})$ as well as the nominal stress $\underline{N}(\vec{r})$ are not altered by the transformation (at the corresponding points $\vec{r} \rightarrow \vec{r}/a$):

$$\underline{F}^*(\vec{r}) = \frac{\partial \vec{U}^*(\vec{r})}{\partial \vec{r}} + \underline{I} = a \cdot \frac{\partial \vec{U}(\vec{r}/a)}{\partial \vec{r}} + \underline{I} = \underline{F}(\vec{r}/a) \tag{2}$$

$$\longrightarrow W^*(\vec{r}) = W(\vec{r}/a) \tag{3}$$

$$\longrightarrow \underline{N}^*(\vec{r}) = \underline{N}^*(\vec{r}/a) \longrightarrow \mathrm{div}(\underline{N}^*(\vec{r})) = \frac{1}{a} \cdot \mathrm{div}(\underline{N}^*(\vec{r}/a)) \ , \tag{4}$$

where $\underline{I}$ denotes the unit tensor. Consequently, the equilibrium equation is fulfilled if the body force is divided by the scaling factor $a$:

$$\rho_0 \vec{b}^*(\vec{r}) + \mathrm{div}(\underline{N}^*(\vec{r})) = 0 = \frac{1}{a} \cdot \Big( \rho_0 \vec{b}^*(\vec{r}/a) + \mathrm{div}(\underline{N}^*(\vec{r}/a)) \Big) \tag{5}$$

We next consider an infinite continuous body with a spherical hole of radius $R$ at the origin. As a boundary condition, we assume that the spherical inclusion has decreased its radius by $\Delta R$, and we denote the displacement field $\vec{U}(\vec{r})$ as the equilibrium solution. The total strain energy needed for the inclusion to contract (or dilate) can be determined by integrating the strain energy density:

$$E(\Delta R) = \int_R^\infty W(\vec{r}) d^3\vec{r} \tag{6}$$

If we now assume that a spherical inclusion with a radius $a \cdot R$ contracts by $a \cdot \Delta R$, we can use the scaling laws noted above to relate the strain energy to that of the un-scaled contracting inclusion:

$$E^* = \int_{a \cdot R}^\infty W(\vec{r}/a) d^3\vec{r} = a^3 \cdot \int_R^\infty W(\vec{r}) d^3\vec{r} = a^3 \cdot E \tag{7}$$

In equilibrium, the strain energy of the contracted inclusion depends only on $R$, $\Delta R$, and the scaling factor $a$:

$$E(a \cdot R, a \cdot \Delta R) = a^3 \cdot E(R, \Delta R) \tag{8}$$

Finally, we show that the normal surface pressure $P$ induced by the contraction of the spherical inclusion only depends on the relative contraction $\Delta R/R$, but not on the scaling factor $a$:

$$\frac{\partial E(R,\Delta R)}{\partial \Delta R} \cdot \frac{1}{4\pi R^2} = \frac{\frac{\partial E(a\cdot R, a\cdot\Delta R)}{a^3}}{\frac{\partial(a\cdot\Delta R)}{a}} \cdot \frac{1}{4\pi R^2} = \frac{\partial E(a\cdot R, a\cdot\Delta R)}{\partial(a\cdot\Delta R)} \cdot \frac{1}{4\pi a^2 R^2} = P(\Delta R/R) \tag{9}$$

Given a fixed surface pressure, a simulated displacement field $\vec{U}(\vec{r})$ of a spherical inclusion with radius $R$ is thus directly related to the deformation field $\vec{U}^*(\vec{r})$ of a spherical inclusion with radius $R^*$ by proper re-scaling with the factor $a = R^*/R$: $\vec{U}^*(a\cdot\vec{r}) = a\cdot\vec{U}(\vec{r})$.


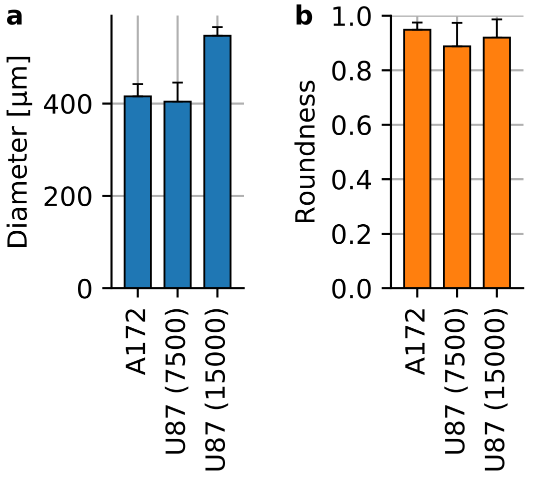


**Supplementary Fig. 1. Spheroid shape and size. a:** Spheroid diameter for A172 containing 15,000 cells at the time of seeding (n=16) and U87 spheroids with 7,500 cells (n=17) and with 15,000 cells (n=15). **b:** Spheroid roundness measured as $4\cdot\text{Area}/(\pi\cdot\text{MajorAxis}^2)$. Error bars denote 1 st.dev.

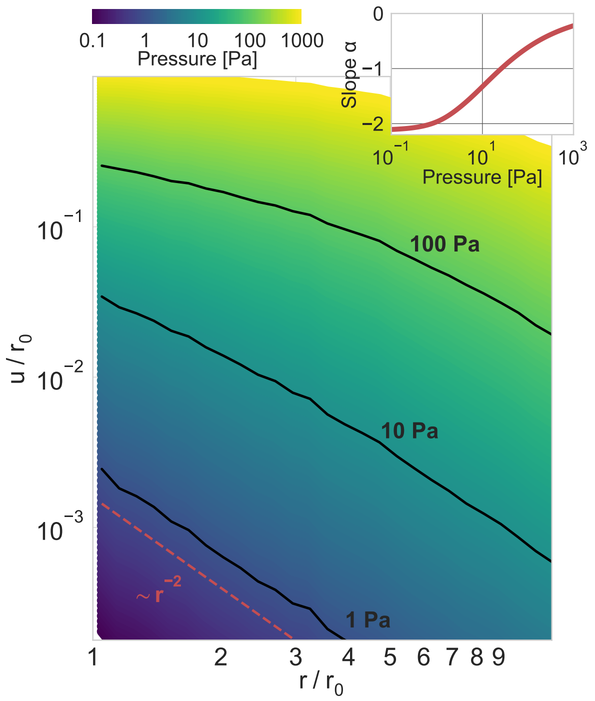


**Supplementary Fig. 2.** **Power-law scaling of deformation fields.** Normalized absolute deformations as a function of the normalized distance, for material simulations with an inbound pressure on the surface of a spherical inclusion ranging from 0.1 Pa to 1000 Pa. The inset shows the power-law exponent $\alpha$ of the deformation field as a function of the inbound pressure (for the near field, $r/r_0 < 2$), illustrating the long-range force transmission in collagen due to strain stiffening.

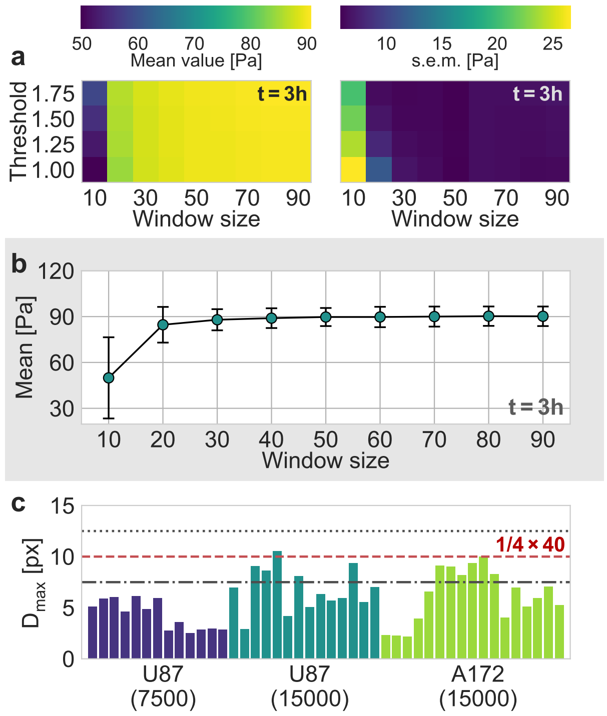


**Supplementary Fig. 3.** **Evaluation of Particle Image Velocimetry. a:** Estimated mean pressure value (left) and the corresponding standard error (right) of an exemplary U87 spheroid, 3 h after the experiment started, as a function of the window size and the signal-to-noise threshold used in the PIV-analysis. **b:** Mean estimated pressure as a function of the window size, for a signal-to-noise threshold of 1.0, i.e. without any filtering of the estimated deformations. **c:** The 99$^{th}$ percentile of all measured absolute displacements during the first two hours of the experiment, $D_{max}$, for all individual spheroids. The dashed red line represents the upper boundary of $D_{max}$ for a window size of 40 px, according to the one-quarter-rule. The dot-dashed line and the dotted line correspond to the upper boundary for a window size of 30 px and 50 px, respectively.

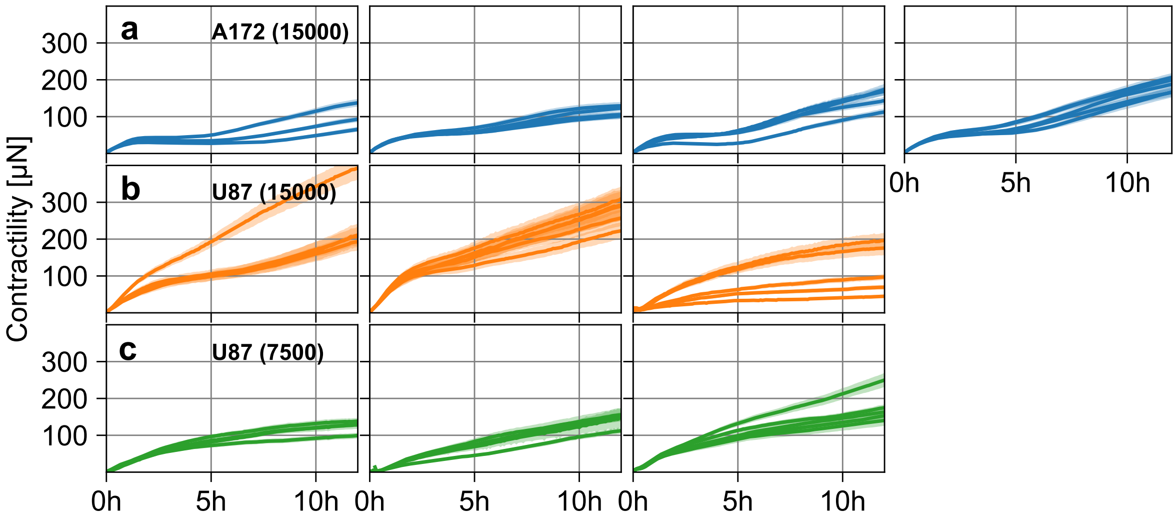


**Supplementary Fig. 4.** **Spheroid contractility over time. a:** Estimated pressure (blue lines) and corresponding standard error (blue shading) of individual A172 tumor spheroids containing 15,000 cells as a function of time, for four separate experiments (left to right). **b:** Same as in (a), for three experiments with U87 tumor spheroids containing 15,000 cells. **c:** Same as in (a), for three experiments with U87 tumor spheroids containing 7,500 cells.

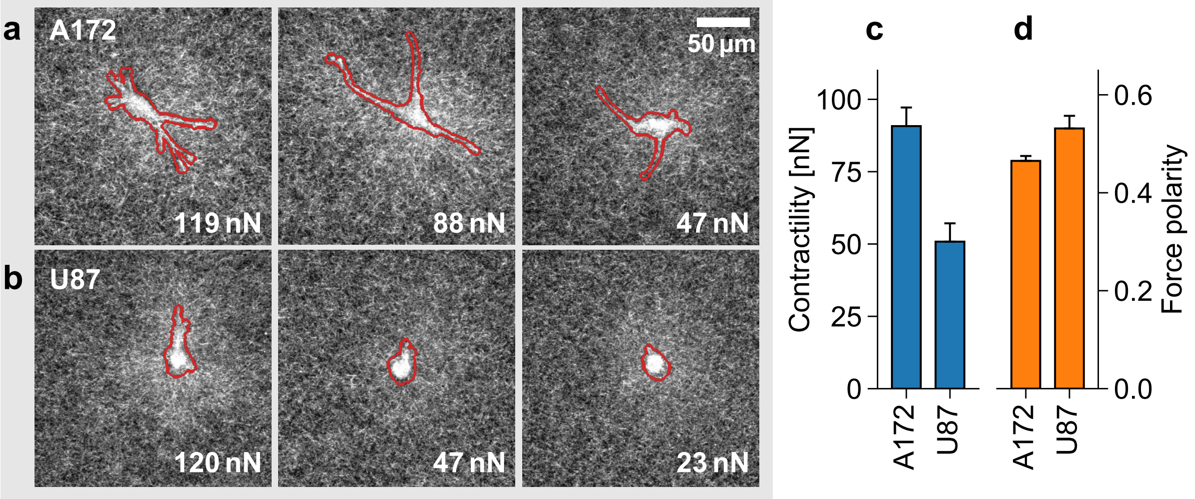


**Supplementary Fig. 5.** **Single-cell 3D traction force microscopy. a:** Exemplary confocal reflection images of A172 glioblastoma cells and the surrounding collagen fibers. The images show the summed pixel intensities along the z-axis, within ±12.5 µm of the z-position of the individual cells. Cell outlines are determined from the transmission channel intensities and are indicated in red. **b:** Same as in (a), but for U87 glioblastoma cells. **c:** Median cell contractility as measured by 3D traction force microscopy (A172: n=90; U87: n=86). d: Median polarity of the force field around individual glioblastoma cells. Error bars denote 1 standard error of the median.
